# ScaR - A tool for sensitive detection of known fusion transcripts: Establishing prevalence of fusions in testicular germ cell tumors

**DOI:** 10.1101/518316

**Authors:** Sen Zhao, Andreas M. Hoff, Rolf I. Skotheim

## Abstract

Bioinformatics tools for fusion transcript detection from RNA-sequencing data are in general developed for identification of novel fusions, which demands a high number of supporting reads and strict filters to avoid false discoveries. As our knowledge of bona-fide fusion genes becomes more saturated, there is a need to establish their prevalence with high sensitivity. We present ScaR, a tool that uses a scaffold realignment approach for sensitive fusion detection in RNA-seq data. ScaR detects a set of 50 synthetic fusion transcripts from simulated data at a higher sensitivity compared to established fusion finders. Applied to fusion transcripts potentially involved in testicular germ cell tumors (TGCTs), ScaR detects the fusions *RCC1-ABHD12B* and *CLEC6A-CLEC4D* in 9% and 28% of 150 TGCTs, respectively. The fusions were not detected in any of 198 normal testis tissues. Thus, we demonstrate high prevalence of *RCC1-ABHD12B* and *CLEC6A-CLEC4D* in TGCTs, and their cancer specific features. Further, we find that *RCC1-ABHD12B* and *CLEC6A-CLEC4D* are predominantly expressed in the seminoma and embryonal carcinoma histological subtypes of TGCTs, respectively. In conclusion, ScaR is useful for establishing the frequency of known fusion transcripts in larger data sets and detecting clinically relevant fusion transcripts with high sensitivity.

**Availability:** https://github.com/senzhaocode/ScaR

## Introduction

Fusion genes and fusion transcripts are important in cancer biology and are often entirely cancer specific, making them attractive as biomarkers. Their attention started with the discovery of the Philadelphia chromosome and the resulting *BCR-ABL1* fusion in patients with chronic myelogenous leukemia (CML) (1–4). In the 1980s and 90s, multiple recurrent fusions were discovered and characterized with chromosome banding and fluorescence in situ hybridization (FISH). These techniques were biased towards detection of fusion genes in hematological cancers and fusions arising from interchromosomal rearrangements (5). With the advent of high-throughput parallel RNA sequencing (RNA-seq) technology, the nomination rate of novel fusion transcripts in both hematological and solid tumor types has exploded. This is underlined with 20731 fusion transcripts being detected in 9966 cancer samples (33 cancer types) from The Cancer Genome Atlas (TCGA) consortium alone (6). Importantly, 83% of these fusion transcripts are detected in single cancer samples and are thus not recurrent. This statistic underlines that fusion transcripts are commonly expressed in cancer, often as a result of increased genomic instability, and that only a minority of these are selected for and act as oncogenic drivers. Therefore, to minimize the detection of additional non-recurrent or nomination of even non-existing (false positive) fusion transcripts, most available fusion finder tools have focused on maximizing specificity.

Nevertheless, several recurrent fusion genes have been indicated as targetable molecular alterations in personalized cancer medicine. These includes for example fusion genes involving the kinase-encoding genes *ALK* and *ROS1* in non-small cell lung cancer, *BCR-ABL1* in CML, and *NTRK1, FGFR3* and *BRAF* in various cancer types (7). In fact, the FDA recently approved Vitrakvi (larotrectinib) as the second tumor-agnostic pan-cancer drug approved for patients harboring *NTRK* gene fusions without a known acquired resistance mutation, are metastatic or where surgical resection is likely to result in severe morbidity and have no satisfactory alternative treatments (8). In addition, highly cancer specific fusion transcripts have potential as biomarkers for disease detection, monitoring and predicting treatment response.

The ability to detect these fusions at high sensitivity is therefore paramount. This will enable us to determine true prevalence of known fusion transcripts in cohorts of cancer patients, where existing fusion finder tools would provide underestimates in efforts of avoiding the scoring of false positives. However, since we in those cases are not searching for novel fusions, we are not risking much by lowering the specificity demands. In more detail, much effort has been invested in developing approaches for fusion transcript detection from RNA-seq data, and a long list of different tools have been developed for this task (Table S1). The performance of fusion finder tools has been shown to vary according to the data set to which they are applied, and none achieve a perfect sensitivity (9, 10). Most of the currently available tools use similar approaches to align reads, nominate fusion transcript breakpoints *de novo* based on supporting reads or read pairs and apply various filters to reduce noise from artifact fusion transcript sequences or the presence of chimeric transcripts in normal cells. A few of the tools available have an option to take a user provided list of known fusion genes and works to force the nominated fusion breakpoints through the list of strict filters *(e.g.* the --focus parameter in FusionCatcher). However, none of them provide a way to directly assess specific fusion breakpoints with high sensitivity. This is underlined in a case where a simple search of a chimeric sequence in raw sequencing data, using the unix tool *“grep”* outperformed the sensitivity of several established fusion finder tools (11). As the knowledge of fusion transcripts and their clinical impact expands together with an increasing amount of patients with RNA-seq data available, there is therefore a need for a tool that can establish the presence of already known and validated fusion transcripts in RNA-seq data with superior sensitivity.

A type of cancer for which no recurrent fusion genes have been established as biomarkers or drug targets is testicular germ cell tumors (TGCT), which is the most commonly diagnosed cancer among young men (12). In fact, not much effort has been done to introduce genomics based personalised medicine for this disease. Although TGCT patients have among the highest survival rates, the treatment choices are few, and side effects are often profound. Further, since the patients are young, serious side effects may affect many decades of their life (13). Therefore, research on fusion genes as potential biomarkers or therapeutic targets in TGCT is of priority. We recently described the detection and characterization of recurrent fusion genes in TGCT (14). TGCT is a disease with distinct histological subtypes including seminomas and nonseminomas, where the latter can be subdivided into pluripotent embryonal carcinomas and more differentiated subtypes; teratomas, yolk sac tumors, and choriocarcinomas. The pluripotent phenotype of malignant TGCTs has similarities to that of embryonic stem cells (15). Studying these cancers can therefore shed light on cancer biology in a context of pluripotency. We previously also showed that the expression of the fusion transcripts *RCC1-ABHD12B* and *RCC1-HENMT1* is reduced upon *in vitro* differentiation of the EC cell line NTERA2 (14). It is therefore of interest to explore the frequency and distribution of the, sometimes weak, expression of these previously identified fusion transcripts in larger cohorts of TGCTs.

Based on the identified need for a sensitive approach to evaluate the recurrence of known fusion transcripts, we herein report the development of a new tool ScaR - **Sca**ffold **R**ealignment. We present benchmarking of ScaR on simulated data and apply it to investigate the prevalence of previously identified fusion transcripts in an extended cohort of TGCTs.

## Materials and Methods

### RNA-sequencing data

We downloaded and processed paired-end RNA-seq raw fastq files of 150 TGCT samples from The Cancer Genome Atlas (TCGA) project (dbGAP accession: phs000178.v9.p8)(16). There were a median of 58.3 million pairs of reads per sample (min: 27.3 million - max: 107.3 million) with read length of 48 × 2 bp (see Table S2 for detailed RNA-seq metrics and sample information). We further downloaded and processed paired-end RNA-seq raw fastq files of 198 normal testicular tissue samples of deceased individuals included in the Genotype-Tissue Expression (GTEx) project (dbGAP accession: phs000424.v6.p1)(17, 18). All GTEx tissue samples were taken from healthy testis and the cause of death of individuals is not related to cancer, according to clinical data from GTEx (Table S4). There were a median of 42.8 million pairs of reads (min: 27.8 million - max: 132.2 million) with read length of 76 × 2 bp (Table S2). Paired-end RNA-seq data from the ES cell line Shef3, as described in Hoff *et al.* (14), was used together with simulated RNA-seq data from synthetic fusion transcripts for benchmarking (see Benchmarking and data simulation).

### *ScaR* and the scaffold realignment approach

The main purpose of Scaffold realignment is to evaluate the presence of known fusion transcripts with breakpoint sites at exon boundaries or within exon regions (Figure 1). Scaffold realignment seeks two types of sequence reads to support fusion transcripts, split reads (a read mapping directly across the fusion transcript breakpoint sequence) and spanning reads *(i.e.* the paired reads map to one fusion partner gene each). Split reads are divided into two categories (Figure 1): discordant-split reads *(i.e.* the other read of the pair maps to the fusion gene partner A / B, or across the fusion transcript breakpoint sequence) or singleton-split reads *(i.e.* the other read of the pair does not map to the transcriptome or genome). The pipeline is divided into four steps: (i) build reference sequences (scaffolds), (ii) read alignment to reference sequences, (iii) read re-alignment to genome sequences and (iv) summarize split read alignments across samples.

**Figure 1:**
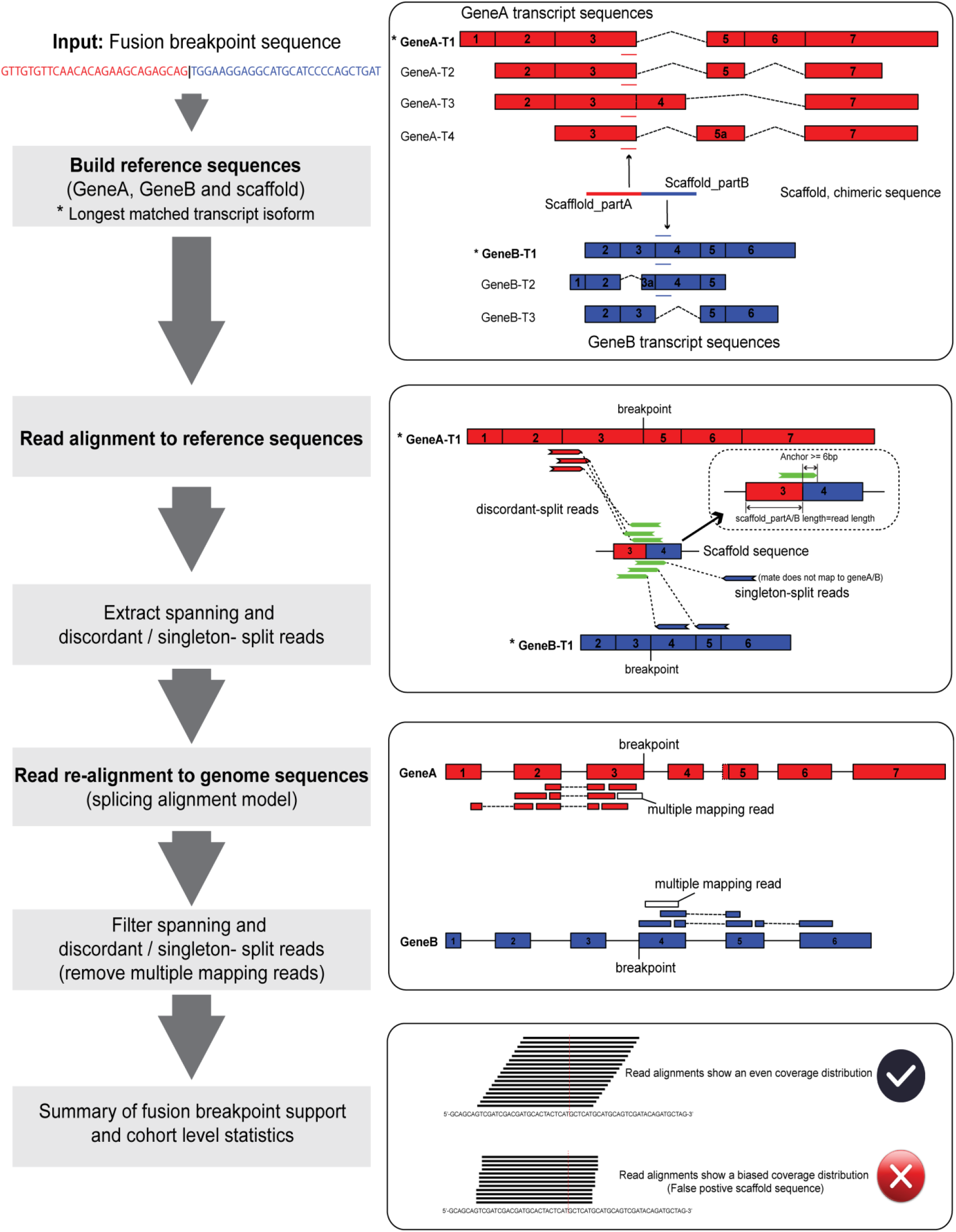
Overview of the scaffold alignment approach – ScaR

#### Build reference sequences

In the first step, a given breakpoint sequence supporting a fusion transcript is split in two sequences at the breakpoint site, which corresponds to Scaffold_partA and Scaffold_partB (Figure 1). If the sequences are longer than the read length used in the sequencing experiment, they are trimmed to match the length of the reads. Each sequence is then screened against all cDNA sequences of gene partners to match parental transcripts (3 transcriptome assembly annotations are packaged with the tool: Ensembl release 89 as default, and GENCODE release 27 or UCSC annotation based on GenBank release 225 and RefSeq release 86 as optional). ScaR also allows user-provided reference annotations if the breakpoint sequences are not previously annotated in the three transcriptome resources. If the sequence match more than one transcript, the longest one *(e.g.* GeneA-T1 and GeneB-T1 marked as * in Figure 1) is selected to represent the gene. If the sequences are shorter than read length, the sequences are extended from the 5’-end of the matching transcript of Scaffold_partA and the 3’-end of the matching transcript of Scaffold_partB, respectively to match the read length. The extended sequences are re-assembled to a new breakpoint sequence scaffold, which together with the sequences of the targeted transcripts from gene A / B serves as a reference sequence for read alignment.

#### Read alignment to reference sequences

To detect the presence of reads supporting the fusion breakpoint, paired-end reads are aligned to the custom scaffold reference using HISAT2 (19). Briefly, an index of the custom scaffold reference sequence is built using *hisat2-build* with default parameters. Paired-end reads are then aligned to the reference sequences using –no-spliced-alignment model with --no-softclip setting. On the basis of the aligned SAM/BAM files, we retrieve three types of mapping reads: discordant-split reads, singleton-split reads and spanning reads. To increase mapping specificity, a minimum anchor length of 6 bp is required (by default) for split reads that map to the fusion breakpoint sequence (Figure 1). All supporting read-pairs of these three mapping types are extracted and saved as fastq files.

#### Read re-alignment to genome sequences

The supporting reads of a fusion breakpoint are further evaluated at a genomic level by aligning all extracted reads to the human reference genome (GRCH38) using HISAT2 -- spliced-alignment model with -no-softclip setting (Figure 1). Supporting reads that are found to align to multiple locations are filtered out. This approach improves specificity and ensures that supporting reads that originate from repetitive sequences or gene homologs are not included in the support of a fusion breakpoint. In this step, singleton-split reads are also renamed as discordant-split reads if the unmapped read could be aligned uniquely to a gene partner at the genomic level.

#### Summary of fusion breakpoint support and cohort level statistics

A minimum support of two discordant-split reads are required to call a positive fusion breakpoint in a given sample. In addition, when the coverage of the fusion transcript is low, supported by only two or three split reads for each sample, the read coverage can show an uneven distribution between Scaffold_partA or Scaffold_partB regions. This uneven distribution can be attributed to either a sampling bias of a random distribution, or an indication of artifact fusion sequences. For a better overview of the mapping distribution for a given scaffold, split reads across all samples in a cohort can be concatenated and aligned to the scaffold sequence. A Fisher’s exact test is then applied to test whether there is a significant bias in the distribution of the number of reads mapped to the upstream and downstream parts of the fusion scaffold sequence (Figure 1).

### Benchmarking and data simulation

To compare the performance of our scaffold alignment approach to that of established *de novo* fusion finders, we applied it together with deFuse v.0.7.0 (20) and FusionCatcher v.0.99.5a (21) on the external TGCT and normal testis data sets from TCGA and GTEx. We searched for the fusion transcripts *RCC1-ABHD12B, RCC1-HENMT1, CLEC6A-CLEC4D and EPT1-GUCY1A3*, as previously identified and characterized in TGCT by Hoff *et al.* (14). The Unix command line tool *grep* was also applied as a simple blunt tool for comparison to our scaffold approach. A string of 15 bp matching the gene on each side of the known breakpoints (30 bp total) were searched for in the fastq files using *grep* and a minimum of two split reads were required for a positive call.

To further benchmark our scaffold alignment approach on a controlled data set, we simulated RNA-seq reads from synthetic fusion transcripts using the MAQ v0.7.1 tool (22). Briefly, we simulated paired-end reads from in total 50 synthetic fusion transcripts. Here, the fusion transcripts were generated between random partner genes that were not paralogs and also with an intergenic distance of more than 50 kb. The minimum combined length of synthetic fusion transcripts was set to 500 bp, with a minimum upstream and downstream sequence length of 100 bp. Paired-end reads (76 bp each) with settings of background mutation rate, -r = 0.0001, fraction of indels, -R = 0.01 and a insert size of 170 bp (SD=25 bp) were simulated. Further, different amounts of synthetic reads to match a gradient sequencing depth of the synthetic fusion transcripts (5X, 10X, 20X, 30X, 50X, 80X, 100X, 150X and 200X) were generated (See Table S3) and then mixed with the RNA-seq reads from the embryonic stem cell line Shef3 (background data). DeFuse, FusionCatcher and *grep* were then applied on this synthetic data set, with identical settings to that previously described. For the scaffold alignment approach, we generated scaffolds of the 50 synthetic transcripts and required a minimum of two discordant split reads as support. The sensitivity of these tools to detect the synthetic fusion transcripts in different mixtures were compared and reported.

### TGCT hierarchical clustering and differential expression analysis

To perform hierarchical clustering and differential expression analysis of the 150 TGCT samples from the TCGA cohort we acquired raw gene count data produced by HTSeq-count from NCI’s Genomic Data Commons (http://xena.ucsc.edu) as well as clinical data including the International Classification of Diseases for Oncology (ICD-O) morphological codes (the latter being available for 134 of the 150 samples; Table S4). Mutation data for the 150 samples was also acquired from cBioportal. The DESeq2 R package (23) was used to perform data normalization and differential expression analysis. Genes that were not expressed across the cohort were removed from further analyses. Prior to performing principal component analysis (PCA) and hierarchical clustering, variance stabilizing transformation was applied on the raw counts. PCA was then performed with the top 500 variable genes used for principal components. Hierarchical clustering was performed on the transformed raw counts using the top 50 most variable genes, clustering on both samples and genes. Clustered heatmaps were produced with the pheatmap R package, plotted together with annotation tracks including ICD-O histological subtypes, fusion transcript status (determined by ScaR) and mutation data of known TGCT driver genes. Mutation status was plotted for genes previously implicated in TGCT and that were mutated in two or more samples in the TCGA cohort. Differential expression analysis was performed on *RCC1-ABHD12B* positive samples vs negative samples and *CLEC6A-CLEC4D* positive vs negative samples, both controlling for the effect of ICD-O histology subtypes.

## Results

### Overview of the ScaR workflow

Here, we sought to establish the frequency of known fusion transcripts in a larger cohort of TGCT patients and we report the development of ScaR - a tool for sensitive detection of known fusion transcripts, which is openly available at https://github.com/senzhaocode/ScaR. ScaR takes any fusion scaffold sequence as input together with raw RNA-seq data, to return the number of spanning and discordant- / singleton-split reads supporting the scaffold sequence (Figure 1). Finally ScaR can summarize the number of of supporting reads across a larger cohort. We applied ScaR to investigate the recurrence of four previously described fusion transcripts *(RCC1-ABHD12B, CLEC6A-CLEC4D, RCC1-HENMT1 and EPT1-GUCY1A3)* in 150 primary TGCT samples using RNA-seq data from TCGA. Overall, we find that ScaR has a sensitivity which is superior to *de novo* tools such as deFuse and Fusioncatcher and the basic grep method in detection of four known fusion transcripts in TGCT.

### Optimization of ScaR parameters

To balance sensitivity and specificity for fusion transcript detection with ScaR, we investigated the sensitivity of detecting the TGCT fusion transcripts with a variable threshold. We also applied the *de novo* fusion finder tools, deFuse and FusionCatcher as well as the basic grep method, to provide a reference for the performance of ScaR. As expected, the detection rate decreased with increasing the minimal threshold of required split reads for all four methods, but ScaR consistently achieved a higher sensitivity compared with the other three tools when setting the threshold below 5 required split reads (Figure 2). All of the four tools show a low sensitivity of detection and a high false negative rate when strict criteria (split read number > 5) are applied for fusion nomination. To evaluate the reads mapping to the different scaffold sequences, ScaR has the ability to concatenate all supporting split reads from a given cohort (in this case the 150 TGCT samples) and align them to the scaffold. For example from this cohort, 59 samples have detectable *RCC1-HENMT1* with a threshold set to one split read, but 51 of them have only one discordant-split read support (Figure 2C; Table S5). We found that 62 reads from 38 of the 51 samples aligned to a scaffold sequence that show a biased distribution around the scaffold breakpoint sequence with a shift towards the *RCC1* part of the scaffold (p = 2×10^−12^; Fisher’s exact test; Figure S1J), indicating that these are false positives. The same pattern was observed for the fusion scaffold *RCC1-ABH12B_alt1* (Figure S1B), but without a significant p-value, probably due to the small number of supporting reads. This coverage bias in the consensus of split read alignments indicates that the reads mapping to the breakpoint scaffold sequences of *RCC1-HENMT1_alt1* and *RCC1-ABH12B_alt1* are most likely mapping artifacts and that these fusion scaffolds represent false positives. These are therefore excluded from further analysis. Overall, from these results, we find that a minimal requirement of two discordant-split reads represents a good balance between sensitivity and specificity for fusion nomination by the ScaR approach, which is further used as a threshold for fusion detection in this study.

**Figure 2.**
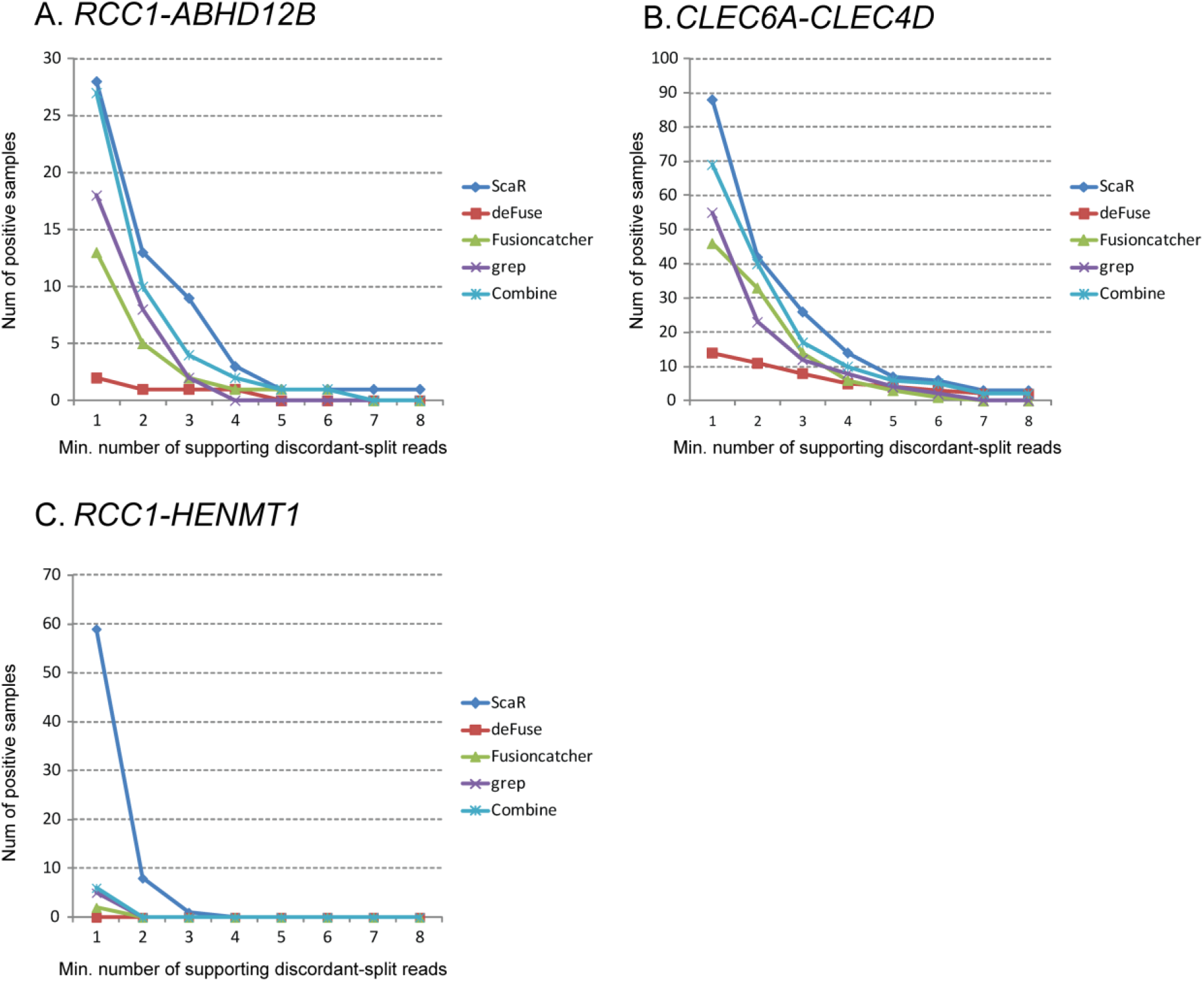
ScaR performance on TGCT data from TCGA. Comparison of sensitivity between ScaR and other tools (deFuse, Fusioncatcher, grep and Combine; combination of the three other tools) for fusion transcripts *RCC1-ABHD12B* (A), *CLEC6A-CLEC4D* (B) and *RCC1-HENMT1* (C) across 150 TCGA TGCT samples. The x-axes show an increasing threshold of minimum required supporting split reads. The y-axes show the number of samples with positive detection.

### ScaR - Benchmarking using simulated fusion transcript read data

To evaluate the performance of ScaR on a controlled data set, we simulated RNA-seq data from 50 synthetic fusion transcripts (Table S3). Various amounts of reads were simulated at 5X to 200X coverage of these synthetic fusion transcripts and mixed *in silico* with real RNA-seq data from the ES cell line Shef3 (48.1 million read pairs; 76 bp x 2). Briefly, 97.5 % of the synthetic reads were found to map to the genome. We further compared the performance of ScaR, deFuse and FusionCatcher to detect these fusion transcripts. ScaR was able to detect 44 out of the 50 fusion transcripts (88 %) at 5X coverage of simulated data, with median of four split reads and one spanning read (Figure 3 and Table S3). In comparison, FusionCatcher and deFuse detected only 23 and 41 of the fusion transcripts at this level, respectively. When increasing the coverage of the synthetic fusion transcripts to 10X and above, all 50 fusion transcripts were detected by ScaR. DeFuse was only able to detect all 50 synthetic fusion transcripts at 200X coverage and FusionCatcher reached a maximum of 45 detectable fusion transcripts.

**Figure 3:**
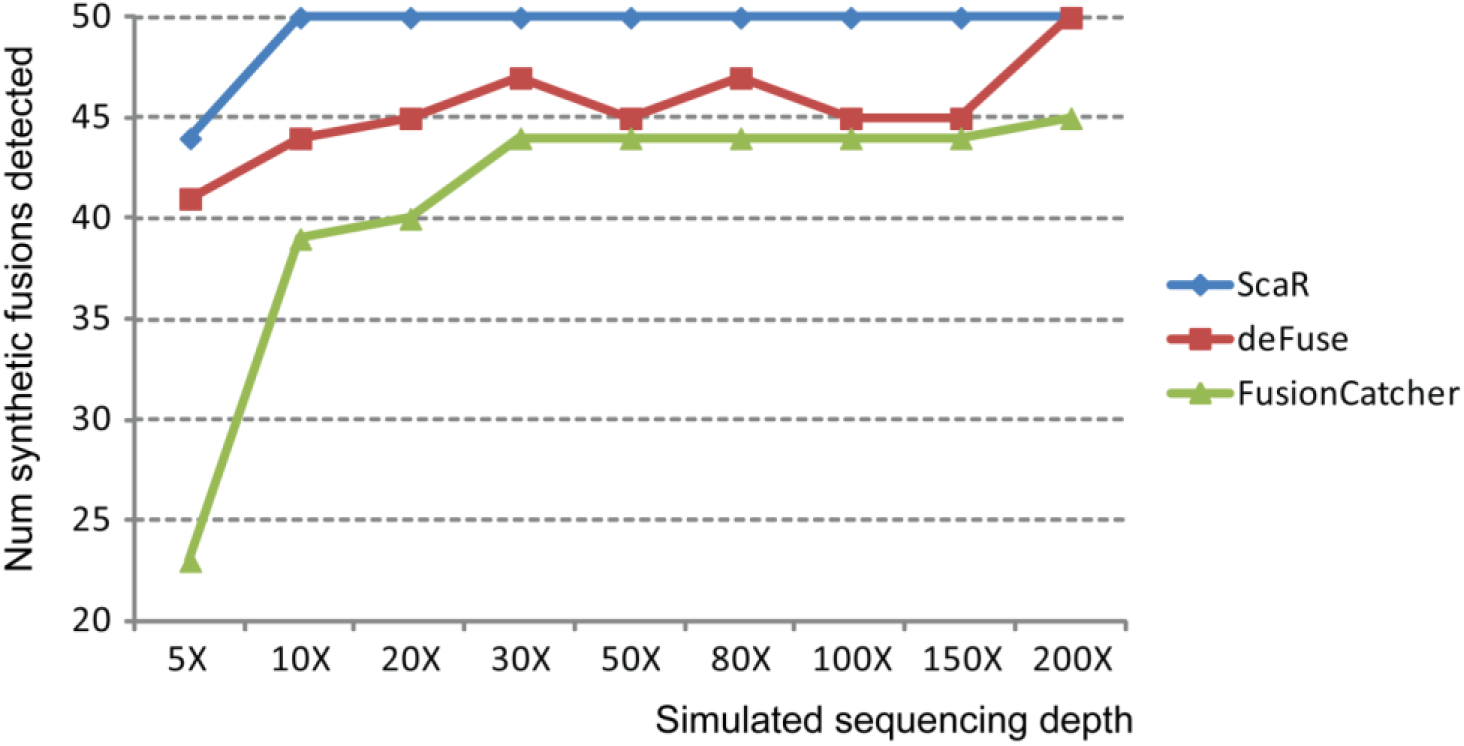
ScaR performance on simulated data. Benchmarking performance of ScaR, deFuse and FusionCatcher on simulated RNA-seq data showing the number of synthetic fusion transcripts detected at simulated coverage levels ranging from 5X to 200X.

### Known fusion transcripts in TGCT are frequently detected by applying ScaR to larger cohorts

To further evaluate the performance of ScaR on real biomedical data we applied ScaR on the TGCT TCGA cohort to detect the previously described fusion transcripts. Specifically, for *RCC1-ABHD12B* (Figure 4A), ScaR detected the fusion transcript in 13 samples (8.7%) with at least two supporting discordant-split reads (Table S5). In comparison, deFuse, FusionCatcher and grep detected the fusion in only one (0.6%), five (3.3%) and eight (5.3%) samples, respectively. By merging the results from these three tools, *RCC1-ABHD12B* was detected in 10 unique samples, where all except one sample (TCGA-XE-A8H4; Figure 4A) overlapped with the positive samples from ScaR. ScaR failed to report the fusion transcript in this sample because one of two supporting split reads is a singleton-split type (Table S5). ScaR detected *RCC1-ABHD12B* in four additional unique samples compared to the other three tools. For *CLEC6A-CLEC4D* (Figure 4B), we evaluated six different fusion breakpoint scaffolds between the two neighbouring genes, as have previously been reported ((14);Table S5). Samples with reads supporting any of these scaffold sequences were regarded as positives. In total, ScaR detected the fusion transcript in 42 (28%) samples, which is higher compared to the frequency identified by deFuse (11; 7.3%), FusionCatcher (33; 22%) and grep (23; 14.7%). Importantly, five of 42 samples detected as positive by ScaR failed to be nominated by any of the three other tools. All positive samples except three cases detected by the deFuse, FusionCatcher or grep are also identified by ScaR. Two of these *(TCGA-2X-A9D6* and *TCGA-WZ-A8D5)* are uniquely identified with grep and have two supporting split reads. For both samples, one of the reads show unspecific multiple alignment at genomic level and is therefore filtered out by ScaR. The third sample *(TTCGA-VF-A8AA)* is exclusively detected by deFuse. We found that the anchor length for supporting split read alignments for this sample is four bp, below the minimum requirement of ScaR. For *RCC1-HENMT1*, ScaR detected the fusion transcript in only one sample when not regarding samples with support for the unreliable *RCC1-HENMT1_alt1* scaffold. DeFuse, Fusioncatcher and grep failed to detect the fusion transcript in any of the 150 samples (Table S5). The fusion *EPT1-GUCY1A3* could not be rediscovered by any of these four tools, with zero spanning and split reads identified. These findings indicate that *EPT1-GUCY1A3* is most likely a private fusion event. We further investigated the scaffold alignments for the samples that were uniquely called by ScaR and not by any of the other tools. For *RCC1-ABHD12B* and *CLEC6A-CLEC4D* which were detected uniquely by ScaR in four and five samples, respectively, we found that the mapping qualities of the reads at the breakpoint sites were of high quality with an even distribution to upstream and downstream regions, suggesting the breakpoints are true positives (Figure 4A and 4B).

**Figure 4.**
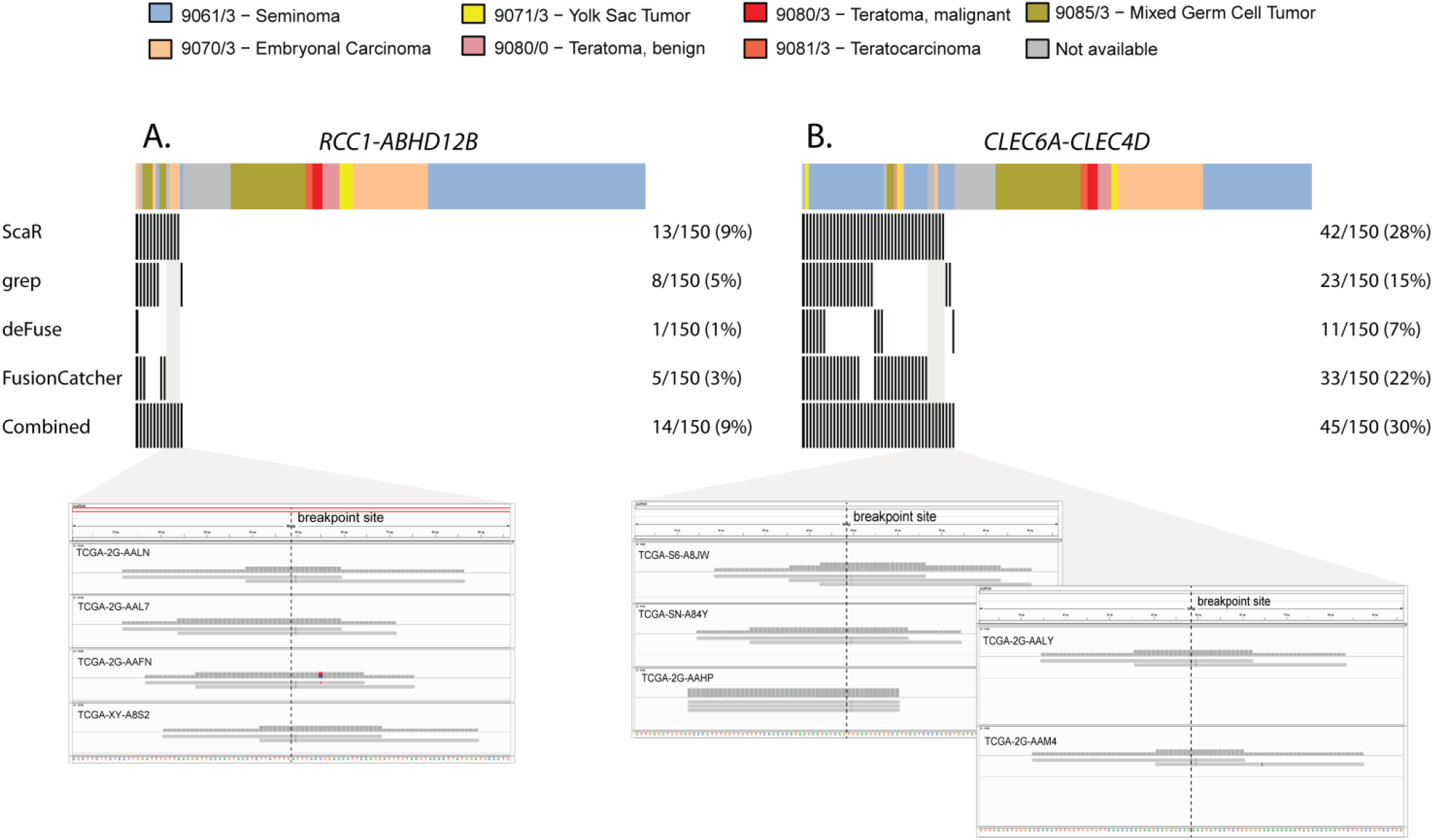
Fusion transcript detection in TGCTs. Overview of TGCT samples from TCGA (n = 150) that are positive for the fusion transcripts *RCC1-ABHD12B* (A) and *CLEC6A-CLEC4D* (B) among the four tools: ScaR, deFuse, Fusioncatcher and Grep. Split read alignments of positive samples uniquely identified by ScaR are visualized using IGV. The ICD-O histology codes are shown as annotated by TCGA.

### TGCT fusion transcripts are malignancy specific and not detected in normal testis tissue samples from the GTEx consortium

We furthermore evaluated the prevalence of these fusion transcripts in 198 normal testicular samples from GTEx project using ScaR. In brief, none of the investigated fusion transcripts could be detected in any of the normal samples (Table S6). For the fusion transcript *CLEC6A-CLEC4D* where the two genes are located only 30 kb apart on chromosome arm 12p, we detected only two split reads and one spanning read all in distinct samples across the 198 samples. Therefore, none of the samples pass the threshold for detection. Similarly, no split or spanning reads are identified for *RCC1-ABHD12B* and one spanning read is identified for *RCC1-HENMT1* across all 198 GTEx samples. Importantly, reads from the GTEx data aligned to genome show a high mapping percentage with a median value of 93.5% (only one sample < 85%) compared to 95.5 % for the TCGA tumor samples. Additionally, the GTEx samples have a median sequencing output of 6.5 Gbp compared to the median sequencing output of the TCGA tumor samples of 5.6 Gbp. These results indicates that the failure to detect the investigated fusion transcripts in normal GTEx samples is not due to differences in sequencing power between the cohorts, and that these fusion transcripts are specifically present in TGCT and not in normal tissue of the testis. This is in accordance with previously published experimental RT-PCR data (14), although then from relatively few samples.

### *CLEC6A-CLEC4D* and *RCC1-ABHD12B* are more frequently detected in the undifferentiated seminoma and embryonal carcinoma like subgroups, respectively

To investigate the biological associations of the frequently identified fusion transcripts *CLEC6A-CLEC4D* and *RCC1-ABHD12B* in data from the TCGA cohort, we performed principal component analysis on gene expression data from the 150 TGCT samples. Not surprisingly, we found that the samples cluster roughly into three groups that correspond well to the annotated ICD-O histological subtypes by TCGA (Figure S2A). The three groups comprise mostly of seminomas, embryonal carcinomas and a third subgroup with the more differentiated histological subtypes and a high frequency of mixed tumors. Further, we performed hierarchical clustering with the 50 most variable genes across the cohort and annotated the samples with somatic mutation calls in known TGCT driver genes, as well as the fusion transcript status, as determined by ScaR (Figure 5). Amongst the top 50 most variable genes we found some of the commonly described stem cell associated genes, such as *NANOG, POU5F1 and SOX2.* Intriguingly, we saw a clear enrichment of *CLEC6A-CLEC4D* expressing samples within the seminoma-like subgroup (p < 0.0001, Fisher’s exact test; Figure 5 and Figure S2C) together with frequent *KIT and KRAS* mutations. For *RCC1-ABHD12B* there was a clear association with the embryonal carcinoma-like subgroup, with 12/13 positive samples clustering within this group (p < 0.0001; Figure 5 and Figure S2B). *CLEC6A-CLEC4D* and *RCC1-ABHD12B* were also largely mutually exclusive, except for in two samples that either had a mixed germ cell tumor or unavailable histological subtype. Further, by differential expression analysis we also found that *RCC1* and *ABHD12B* were significantly upregulated in the *RCC1-ABHD12B* positive subgroup (Figure S3A). Also, both *CLEC6A* and *CLEC4D* were among the highest ranked upregulated genes in the *CLEC6A-CLEC4D* subgroup (Figure S3B).

**Figure 5.**
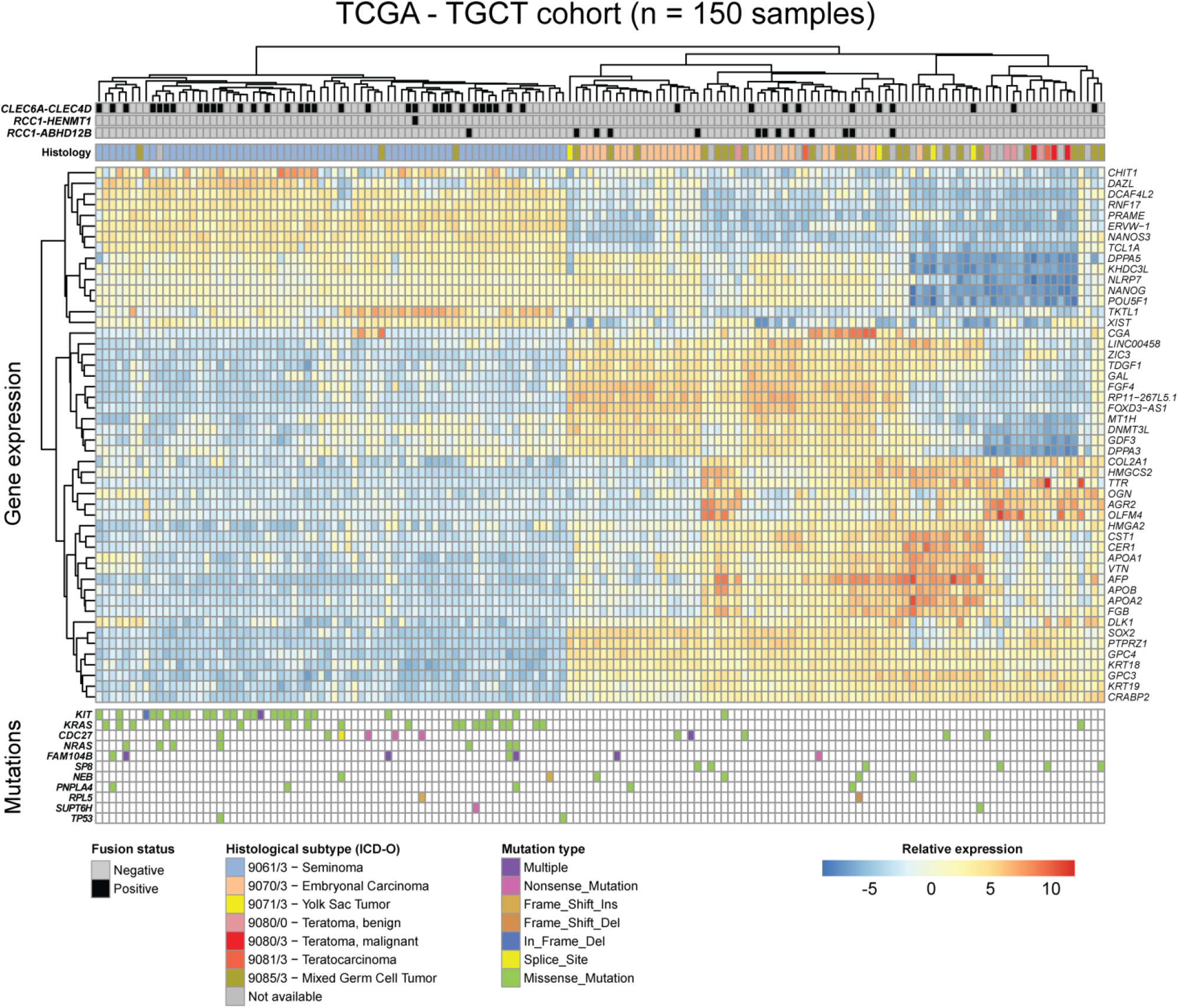
Fusion transcripts in TGCT and associated molecular features. Heatmap showing fusions, somatic mutations, and RNA expression (normalized RNA-seq counts) of the 50 most variable genes across the TCGA cohort. Individual samples are clustered along the horizontal axis while genes are clustered on the vertical axis. Annotation tracks include ICD-O histology codes, and fusion transcript status for *RCC1-ABHD12B, RCC1-HENMT1* and *CLEC6A-CLEC4D*. Somatic mutation status for genes known to be recurrently mutated in TGCT are also shown and colored according to mutation type.

## Discussion

We have developed, tested, and applied the bioinformatics tool ScaR for sensitive detection of known fusion transcripts in RNA-seq data. ScaR efficiently implements a direct scaffold realignment approach, and we have benchmarked the tool on simulated data. Importantly, we have evaluated previously described fusion genes in TGCTs in larger cohorts from the TCGA and GTEx consortia, and demonstrated that ScaR achieves a high sensitivity for known fusions compared to *de novo* fusion finder tools.

The improved sensitivity will be of value as an expanding array of fusion genes with clinical impact are uncovered. Already, multiple fusion genes occurring in cancer are predictive for response to kinase inhibitors, and establishing the presence of such fusion genes and their fusion transcript products in patients prior to treatment is of importance. Improved detection sensitivity for a fusion transcript biomarker can also be important in monitoring a patient’s response to treatment or in detecting minimal residual disease, *e.g.* detecting the presence of *BCR-ABL1* in patients undergoing treatment with the kinase inhibitor *imatinib.* By looking specifically for the fusion transcripts of interest and thereby circumventing the need for strict filters and thresholds to avoid false positives, due to biological and technical noise in RNA-seq data, our approach with ScaR could be better suited for these purposes. Also, as RNA-seq data from more patients and cancer types are becoming available, establishing the prevalence of known fusion transcripts in expanded and new cohorts are of importance. For instance, fusion genes involving the kinases *ALK, RET, ROS1* and *BRAF* have been found in multiple cancer types, expanding the repertoire of cancers that kinase inhibitors could target (24).

We show that our tool has an improved sensitivity compared to the established fusion gene detection tools deFuse and FusionCatcher, both by using simulated data and on real data from TCGA. Although, there are tens of fusion finder tools available, most fusion finder tools build on similar principles by read-alignment, detection of reads or read-pairs that support a fusion breakpoint and applying different filtering criteria. We carried out this comparison using deFuse and FusionCatcher on the basis that deFuse has been an established fusion finder tool for many years (and still maintained) and that FusionCatcher have repeatedly performed well in independent comparison studies on multiple data sets (9, 10, 20). Most *de novo* fusion tools relies on spanning reads to nominate gene partners of fusion genes and split reads are consequently used to refine the exact breakpoint sequences. The amount of spanning read pairs for a given fusion breakpoint is highly dependent on the insert size of the read-pairs in each RNA-seq library. In fact, in sequencing libraries with very short or negative insert sizes (overlapping single-end reads) the number of supporting spanning reads may be very low or completely absent leading to a reduced sensitivity of detection. By providing ScaR with an already known fusion breakpoint we can avoid this bias of insert size. ScaR therefore uses split-reads as the main support for a given fusion breakpoint, but also provides the supporting spanning reads in the output, which may be used for downstream purposes. This is one of the major impacts on the improved sensitivity we see with ScaR compared to *de novo* fusion tools. The unix tool *grep*, which we also compared to, has been used as a direct approach to indicate the presence of fusion transcripts from RNA-seq data (11). However, this approach suffers from requiring a perfect match to the query sequence in RNA-sequencing reads, not allowing for single mismatches, indels or variable anchor lengths. Also, the supporting reads from the *grep* approach are not confirmed to be unambiguously mapping to the breakpoint sequence, or if they potentially map ambiguously to multiple sequences in the genome. ScaR circumvents these drawbacks and improves the sensitivity and specificity compared to the basic grep tool, by using a dedicated aligner for aligning reads to a fusion specific scaffold sequence and further mapping supporting reads back to the genome to avoid ambiguous supporting reads.

ScaR requires the use of a transcriptome annotation and generates the fusion scaffold from exonic sequences of transcripts matched to the input breakpoint sequence. Currently, we include three options of major transcriptome annotation resources (Ensembl, GENCODE and UCSC) in ScaR. In addition, we allow a user-defined annotated reference sequence as input, which could involve non-coding sequences from intronic and intergenic regions. It extends the functionality of ScaR to evaluate fusion transcripts from alternative promoter or new splicing events that are not previously annotated in any of the three major transcriptome annotations.

Here, our aim was to validate the presence and explore on the prevalence of fusion transcripts we previously discovered to be recurrent in a small cohort of TGCTs (14), in a larger cohort from TCGA. Admittedly, we initially found that the frequency of samples positive for these fusion transcripts was much lower than what we previously established with quantitative real-time PCR in our cohort of TGCTs. We therefore explored if these fusion transcripts could be expressed at low levels in a larger number of samples, and that more sensitive approaches were needed to detect this signal in RNA-seq data. By developing and applying ScaR we discovered that these fusion transcripts, especially the read-through *CLEC6A-CLEC4D* and the interchromosomal fusion *RCC1-ABHD12B*, are detectable in a higher frequency of TGCTs than what could be established with *de novo* fusion finder tools. Importantly, we also show that our sensitive detection approach with ScaR does not uncover these fusion transcripts in any samples from a large cohort of normal testis samples (GTEx), indicating that these fusion transcripts, albeit being expressed at low levels, are cancer-specific. Further, by hierarchical clustering on gene expression data from the TCGA, we show that the TGCT samples cluster according to their histological subtypes (16), in line with previous publications on gene expression in TGCT (25). From the heatmap in Figure 5, we see that *CLEC6A-CLEC4D* is significantly enriched in samples of the undifferentiated seminoma-like cluster, while *RCC1-ABHD12B* is significantly enriched in samples of the undifferentiated embryonal carcinoma-like cluster. These findings support our previous results that showed that *RCC1-ABHD12B* expression, but not *CLEC6A-CLEC4D* expression, was significantly reduced when a pluripotent embryonal carcinoma cell line (NTERA2) was differentiated *in vitro* (14). These observations support a biological significance of these fusion transcripts being markers of pluripotent TGCTs.

## Conclusion

We have developed ScaR, a tool that uses a scaffold alignment approach for sensitive detection of known fusion transcripts in RNA-seq data. Such sensitive detection of known fusion transcripts will be of importance in personalized cancer medicine. Further, we have used ScaR to establish that the *RCC1-ABHD12B* and *CLEC6A-CLEC4D* fusion transcripts are frequently detected in TGCTs and associated with the undifferentiated embryonal carcinoma and seminoma histological subtypes.

## Acknowledgments

We would like to thank Jonas M. Strømme for his valuable suggestions and input on the study.

## Author contributions

Conception and design: SZ, AMH and RIS; development and implementation of tool: SZ and AMH; data collection and benchmarking testing: SZ and AMH; data analysis and interpretation: SZ and AMH; writing, review and/or revision of the manuscript: SZ, AMH and RIS; study supervision: RIS

## Supplementary files

**Figure S1**. Split reads across 150 TCGA TGCT samples are concatenated and aligned to the ten chimera scaffold sequences. The number of reads aligned to mostly upstream and downstream regions of the scaffold breakpoint is indicated as “x | y”, where the breakpoint is indicated by a red dotted line, and a P value for the estimate of mapping biases towards either part of the scaffold sequence from a Fisher’s exact test is shown.

**Figure S2**. The first two components from principal components of gene expression data from the 150 TCGA TGCT samples. Data from the top 500 variable genes were used as input. Samples are colored according to: A) histology subtype (ICD-O codes), and detection of the fusion transcripts B) *RCC1-ABHD12B*, C) *CLEC6A-CLEC4D* and D) *RCC1-HENMT1.*

**Figure S3**. Boxplots of log2-transformed normalized count values for A) *RCC1* and *ABHD12B* grouped by *RCC1-ABHD12B* fusion transcript status, as determined by ScaR and B) *CLEC6A* and *CLEC4D* grouped by *CLEC6A-CLEC4D* fusion transcript status. Adjusted p-values from differential expression analysis using DESeq2 and controlling for ICD-histology effect contribution are shown.

**Table S1**. Compiled list of fusion finder tools. The number of citations were extracted from Google Scholar January 2019.

**Table S2**. Total number of paired end reads and mapping statistics from the 150 TGCT TCGA samples and the 198 normal testis samples from GTEx.

**Table S3**. Summary of the RNA-sequencing data simulated by the MAQ tool and used for benchmarking. The 50 fusion genes and their respective gene partners are listed. Further, mapping statistics are shown and results from ScaR, deFuse and FusionCatcher.

**Table S4**. Clinical annotation of the 150 TCGA and 198 GTEx samples.

**Table S5**. Results from deFuse, FusionCatcher, *grep* and ScaR for detecting the fusion transcripts *RCC1-ABHD12B, CLEC6A-CLEC4D, RCC1-HENMT1* and *EPT1-GUCY1A3* in 150 TCGA TGCT samples. The scaffold sequences used for ScaR are also listed.

**Table S6**. Results from ScaR for detecting fusion transcripts *RCC1-ABHD12B, CLEC6A-CLEC4D, RCC1-HENMT1* and *EPT1-GUCY1A3* in 198 normal testis samples from GTEx.

