## Supplementary material for "ScaR - A tool for sensitive detection of known fusion transcripts: Establishing prevalence of fusions in testicular germ cell tumors": Figure S1

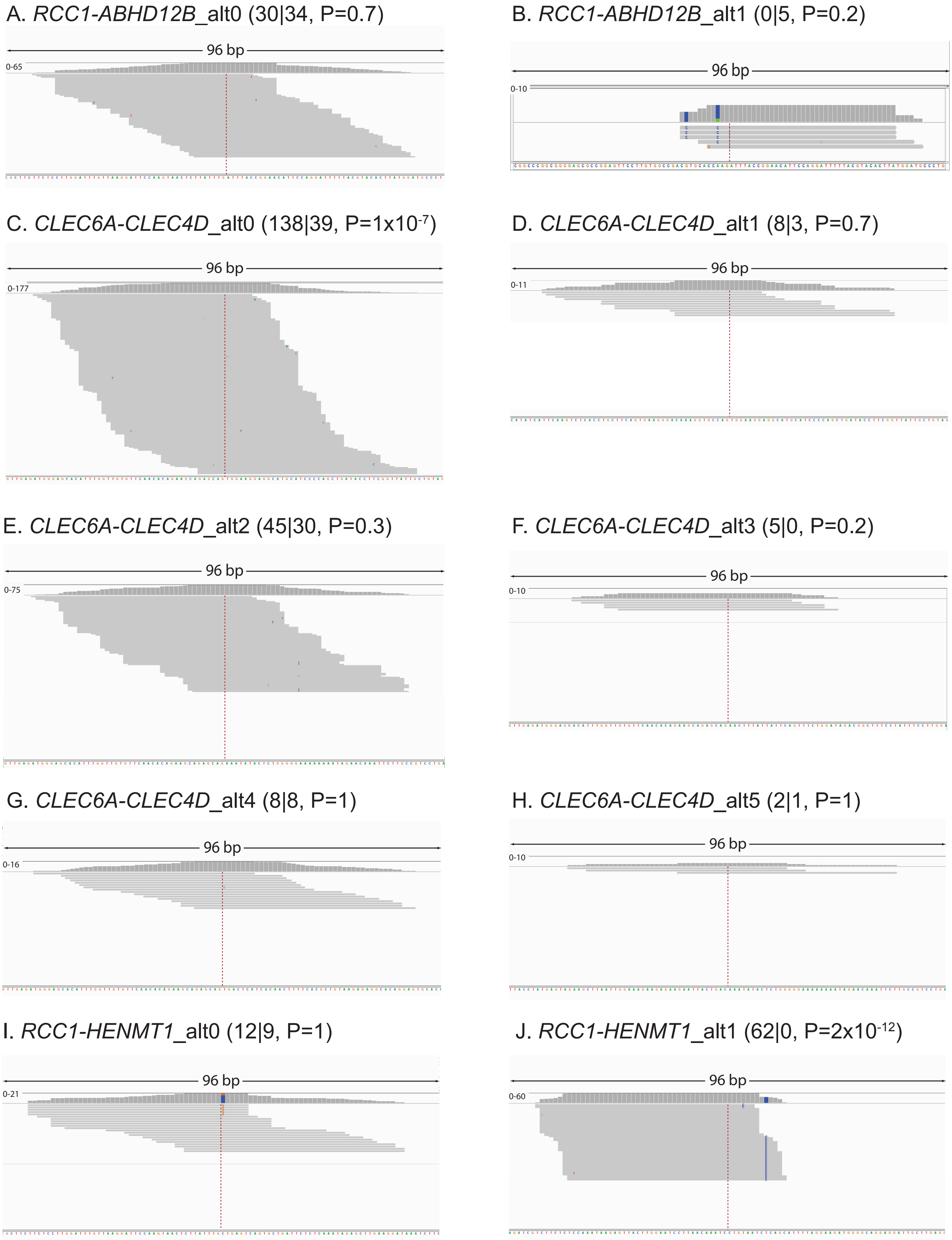


**Figure S1.** Split reads across 150 TCGA TGCT samples are concatenated and aligned to the ten chimera scaffold sequences. The number of reads aligned to mostly upstream and downstream regions of the scaffold breakpoint is indicated as "x | y", where the breakpoint is indicated by a red dotted line, and a P value for the estimate of mapping biases towards either part of the scaffold sequence from a Fisher's exact test is shown.
