## Supplementary material for "ScaR - A tool for sensitive detection of known fusion transcripts: Establishing prevalence of fusions in testicular germ cell tumors": Figure S2

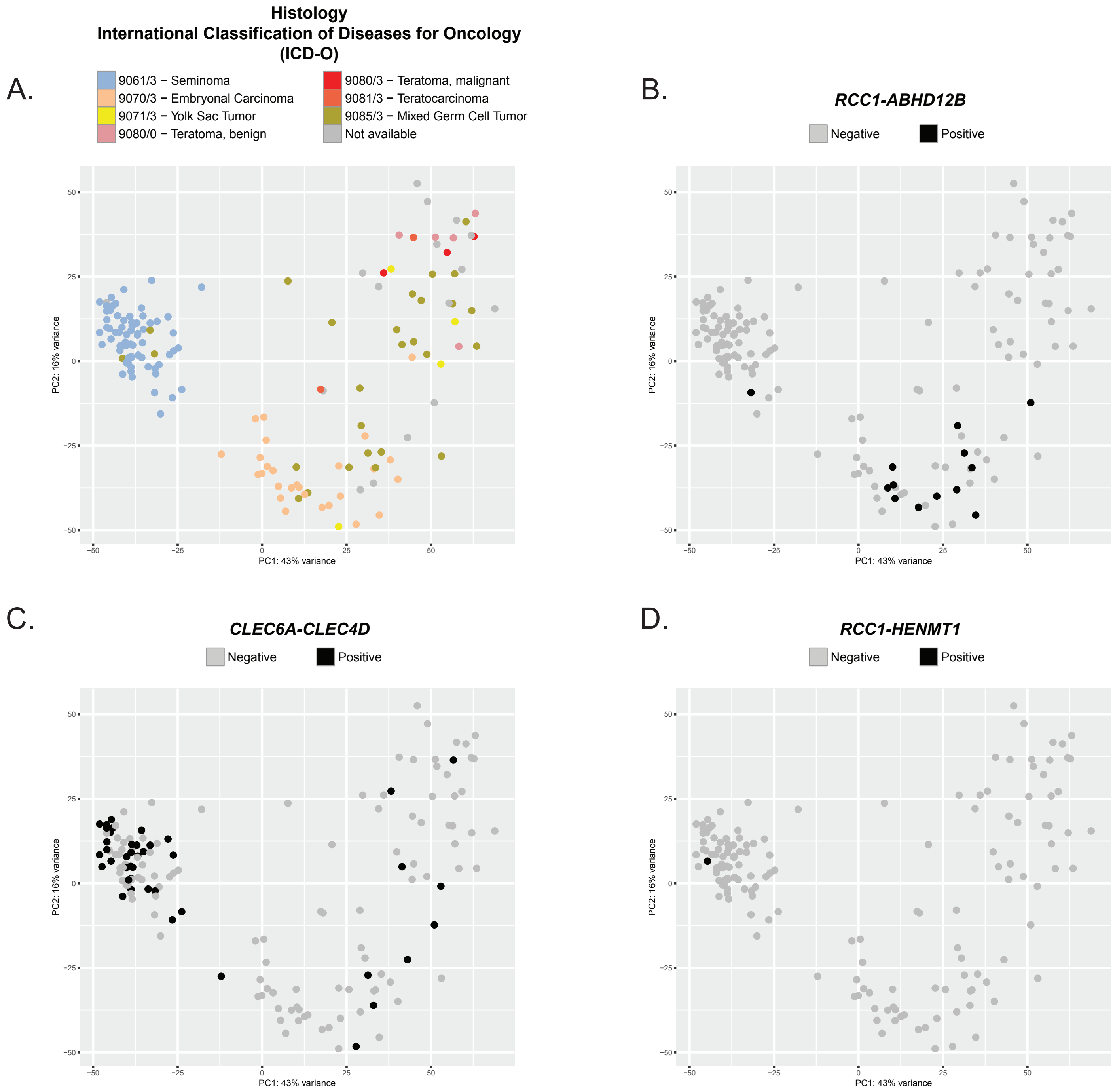


**Figure S2.** The first two components from principal components of gene expression data from the 150 TCGA TGCT samples. Data from the top 500 variable genes were used as input. Samples are colored according to: A) histology subtype (ICD-O codes), and detection of the fusion transcripts B) *RCC1-ABHD12B*, C) *CLEC6A-CLEC4D* and D) *RCC1-HENMT1*.
