## Supplementary material for "ScaR - A tool for sensitive detection of known fusion transcripts: Establishing prevalence of fusions in testicular germ cell tumors": Figure S3

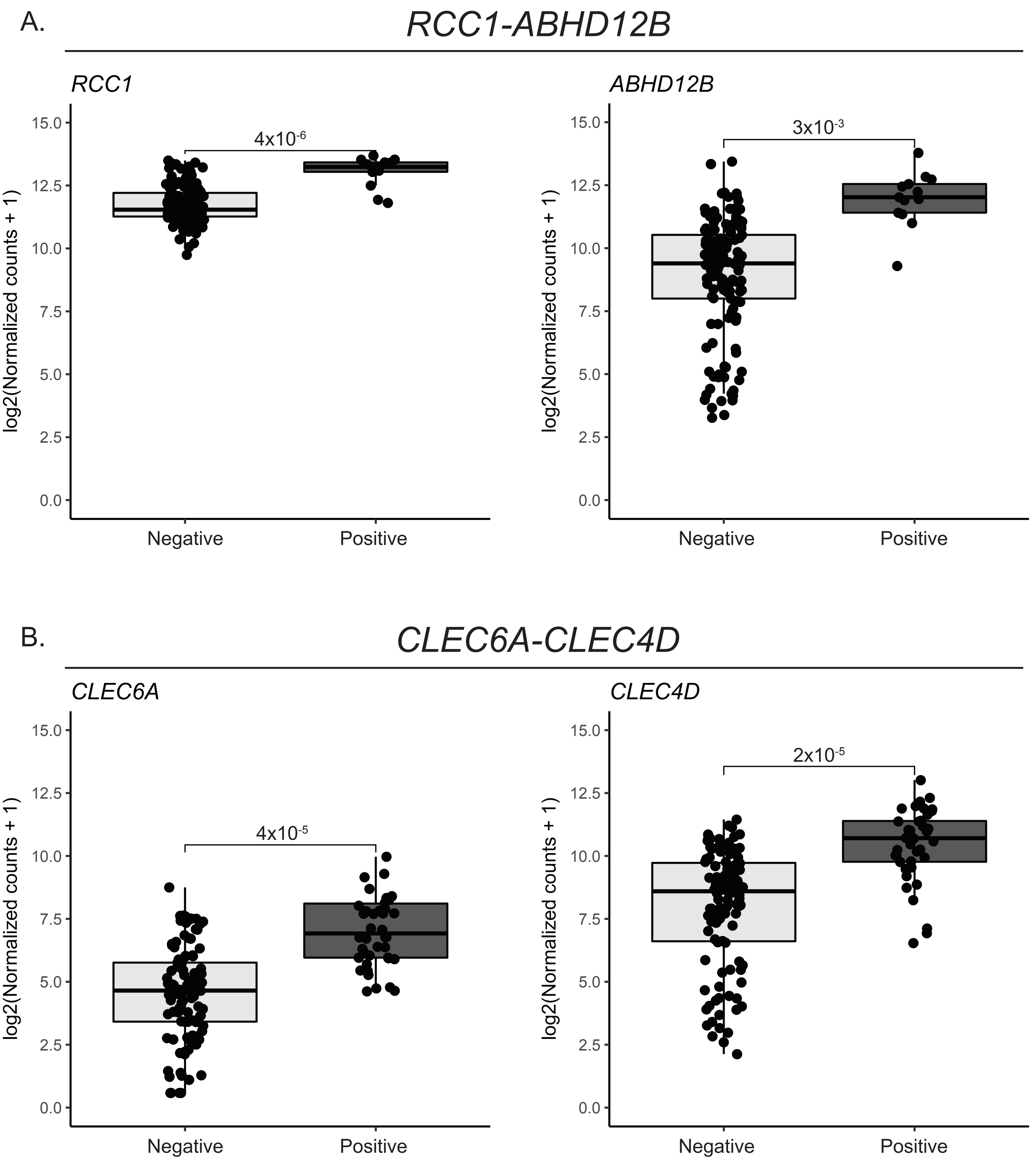


**Figure S3.** Boxplots of log2-transformed normalized count values for A) *RCC1* and *ABHD12B* grouped by *RCC1-ABHD12B* fusion transcript status, as determined by ScaR and B) *CLEC6A* and *CLEC4D* grouped by *CLEC6A-CLEC4D* fusion transcript status. Adjusted p-values from differential expression analysis using DESeq2 and controlling for ICD-histology effect contribution are shown.
